# Comparative study of manganese catalase monomers, interfaces and cage architecture within the ferritin superfamily

**DOI:** 10.1101/2025.06.17.660058

**Authors:** Amrita, Soumyananda Chakraborti, Sucharita Dey

## Abstract

Manganese catalase protein is an example of a protein cage within the ferritin superfamily that focuses on enzymatic catalysis rather than on storage which other ferritin proteins are known for. Formed of 6 homomeric chains, it shows large subunit-subunit sidewise contacts and special interfacial interactions ensuring cage generation with only 6 subunits. We aim to explore manganese catalase at the monomer, subunit-subunit and cage level. As compared to other ferritin subtypes, we found manganese catalase monomers to have greater fraction of β-turns at the secondary structure level and thermophilic manganese catalase monomers to have greater non-polar to polar residue ratio. At the interface level, the placement of subunits in manganese catalase cage with sidewise and angular interface was found to be distinctly different from the parallel and perpendicular interfaces present in other ferritin subtypes. Our study highlights the contribution of sidewise interfaces and terminal extensions in making the cage architecture possible in 6-mer manganese catalases. We also studied the quaternary structure of manganese catalase cage with respect to Classical ferritin (C-ferritin) and found manganese catalase to have smaller cavity volume and cavity surface area than C-ferritin but higher cavity surface to volume ratio. This observation along with the smaller distance between substrate entry point and active site as compared to C-ferritin highlights the structural distinction between catalytic enzymes like manganese catalase and storage proteins like ferritins. Thus, the study gives structural insight into manganese catalase protein cage with focus on its ability to form a cage architecture and show efficient catalytic activity.

## Background

Manganese catalase is an essential enzyme found in a wide range of prokaryotes that neutralizes hydrogen peroxide through catalysis thereby preventing hydrogen peroxide associated cytotoxicity. They are an example of the oxidoreductase category of enzymes where manganese catalase along with other ferritin family of proteins forms a relatively rare cage-like architecture (dihydromethanopterin reductase as only other example). These proteins are quite intriguing due to their unique structural features and potential applications in nanotechnology and biocatalysis. Manganese catalase comprise Mn as a cofactor, which facilitates catalysis of H□O□ into H□O and O□. This property has rendered the protein useful in the medical industry for the treatment of oxidative stress, in the pharmaceutical industry for the removal of peroxide from specific chemical solutions, and in environmental biotechnology for providing oxygen during aerobic bioremediation(Kaushal et al., 2018). Manganese catalases are formed by 6 homomeric chains that self-assemble into a cage architecture with extensive intersubunit contacts and a structural Ca ion cross-linking neighbouring chains(Bihani et al., 2016). Within the manganese catalase family, there exist two distinct structural classes (mesophilic and thermophilic manganese catalases) based on the organisms in which they exist. Thermophilic manganese catalases from *T. thermophilus(Antonyuk et al., 2000)* and mesophilic manganese catalases from *L. plantarum(Barynin et al., 2001)* are the most studied manganese catalases due to the availability of their crystal structures. Thermophilic catalases may have functions other than peroxide decomposition (catalase and peroxidase); meanwhile, mesophilic catalase enzyme exerts a restricted access to H_2_O_2_ and specifically serves as a catalase(Kono and Fridovich, 1983a). Apart from that, these structural classes differ in their monomeric design and the organization of their Mn active site as well as the Ca ion crosslinker which is present in mesophilic catalases but absent in their thermophilic counterpart.

The quaternary architecture of the protein shows Manganese catalase as a Homo-6-mer with a 3-2 point symmetry. Each subunit of the protein is composed of an N-terminus extension, a 4 helix bundle catalytic core characteristic of a Ferritin-like superfamily(Andrews, 2010), and a C-terminus tail. The 2 terminal regions (residues 1-18 and 191-260) distinguish manganese catalases from other ferritin family proteins in having a distinctly extended conformation involving very little secondary structure and wrapping the core like a belt, thereby protecting the core and stabilizing the cage architecture(Barynin et al., 2001). The Manganese catalase catalytic domain comprises a binuclear manganese metal complex consisting of one glutamate carboxylate bridge (μ□,□) between the metal centers. EXXH metal binding motif of the glutamate carboxylate bridge is a common feature of the proteins containing a binuclear metal complex. A separate non-coordinated glutamic acid residue lies above the metal core, forming a H-bond with the solvent molecule attached to Mn ion(Whittaker, 2012). A bulge formed by a pair of Glycine residues in the middle of helix A creates an access channel connecting the catalytic manganese cluster to the solvent-filled core of the protein. The access channel is lined by hydrophilic groups that may restrict the entry of ions while allowing access to neutral species like H□O□, H□O and O□(Barynin et al., 2001).

We try to study the structural and functional characteristics of manganese catalase at the monomer, subunit-subunit interface and cage level, with the aim to explore the similarities and distinctions between the 2 classes of manganese catalases-Mesophilic and Thermophilic manganese catalases as well as between 6-mer manganese catalases and 24-mer Classical ferritins (C-ferritins) in general. The study gives an insight into mechanisms of protein cage assembly at the subunit, interface and cage level and the factors that influence it, thus giving a comparative overview of the protein cages at 2 different length scales(Aumiller et al., 2018).

## Results and Discussion

### Dataset curation

The four of the best characterized(Whittaker, 2012), non-redundant and native non-mutant manganese catalase proteins, PDB:1jku (Barynin et al., 2001) (*Lactobacillus plantarum*), PDB:6j42 (Bihani et al., 2016) (*Nostoc sp.*), PDB: 2v8t (Antonyuk et al., 2000) (*Thermus thermophilus* HB27) and PDB: 2cwl (*Thermus thermophilus* HB8), were taken as the initial reference in the study of manganese catalases. For future studies, these proteins were categorized into mesophilic groups (PDB: 1jku, 6j42) and thermophilic groups (PDB: 2v8t,2cwl) and sequence based analyses were undertaken on their homologs. For a comparative study of manganese catalases with ferritins, the analysis was expanded to native and non-redundant ferritins obtained from our previous dataset - 18 Classical ferritins (C-ferritins), 10 Bacterial ferritins (B-ferritins), 13 Bacterioferritin (Bo-ferritins), 30 Dps (DNA-binding protein from starved cell) (“Physicochemical features of subunit interfaces and their role in self-assembly across the ferritin superfamily,” 2024).

### Sequence and structure-based characterization of the Manganese catalase monomer

Manganese catalase monomer comprises 3 main segments-1) an N-terminal polypeptide 2) a 4-helix bundle catalytic domain and 3) an extended C-terminal tail (Figure 1H and 1I). These long terminal extensions ensure extensive contacts with every other subunit of the protein, ensuring a strong interaction. The 4-helix domain is well conserved across the ferritin superfamily as observed for Classical ferritin (C-ferritin) in Figure 1F. Also every subunit of manganese catalase also constitutes 2 Mn ions responsible for its catalytic activity(Kono and Fridovich, 1983b). Several physicochemical properties were obtained from the Protparam tool(Wilkins et al.,1999) for the Manganese catalase and ferritin monomeric sequences. These properties give us an insight into the structural characteristics of these monomers and the physico-chemical variations between them (Table 1).

**Table 1:** Sequence-based physico-chemical properties of Manganese catalase and Ferritin monomers.

|  | Mesophilic MnCatalase | Thermophilic MnCatalase | C-ferritin | B-ferritin | Bo-ferritin | Dps proteins |
| --- | --- | --- | --- | --- | --- | --- |
| Assembly | 6-mer | 6-mer | 24-mer | 24-mer | 24-mer | 12-mer |
| Number | 31 | 39 | 18 | 10 | 13 | 30 |
| MW | 29846±294 | 33211±371 | 20739±1562 | 20145±1601 | 18925±1080 | 18947±1832 |
| Fraction (Non-polar residues) | 0.47 | 0.54 | 0.45 | 0.47 | 0.47 | 0.48 |
| Fraction (Polar residues) | 0.53 | 0.46 | 0.55 | 0.53 | 0.53 | 0.52 |
| GRAVY | -0.63 | -0.24 | -0.61 | -0.38 | -0.45 | -0.27 |
| Isoelectric Point | 5.03±0.12 | 5.5±0.16 | 5.32±0.36 | 4.99±0.42 | 4.84±0.20 | 5.09±0.44 |
| Helix fraction | 0.68 | 0.65 | 0.9 | 0.89 | 0.85 | 0.84 |
| Turn fraction | 0.27 | 0.28 | 0.1 | 0.1 | 0.1 | 0.11 |
| β-Sheet fraction | 0.05 | 0.07 | 0.0 | 0.01 | 0.05 | 0.04 |
| Top 3 AA | GLY, GLU, LEU | LEU, ALA, GLY | LEU, GLU, ALA | LEU, GLU, ALA | LEU, GLU, ALA | LEU, ALA, GLU |
| Instability Index | 33.6(stable) | 41(stable) | 37.5(stable) | 39.2(stable) | 39.9(stable) | 34.5(stable) |

<MW>:Molecular weight in Daltons
<GRAVY>(Wilkins et al., 1999):The Grand Average of Hydropathy calculated as the sum of hydropathy values of all the amino acids, divided by the number of residues in the sequence.
\* Helix fraction, Turn fraction and β-Sheet fraction data are obtained from the DSSP tool (Kabsch and Sander, 1983) using PDB structures of the proteins.

**Figure 1:**
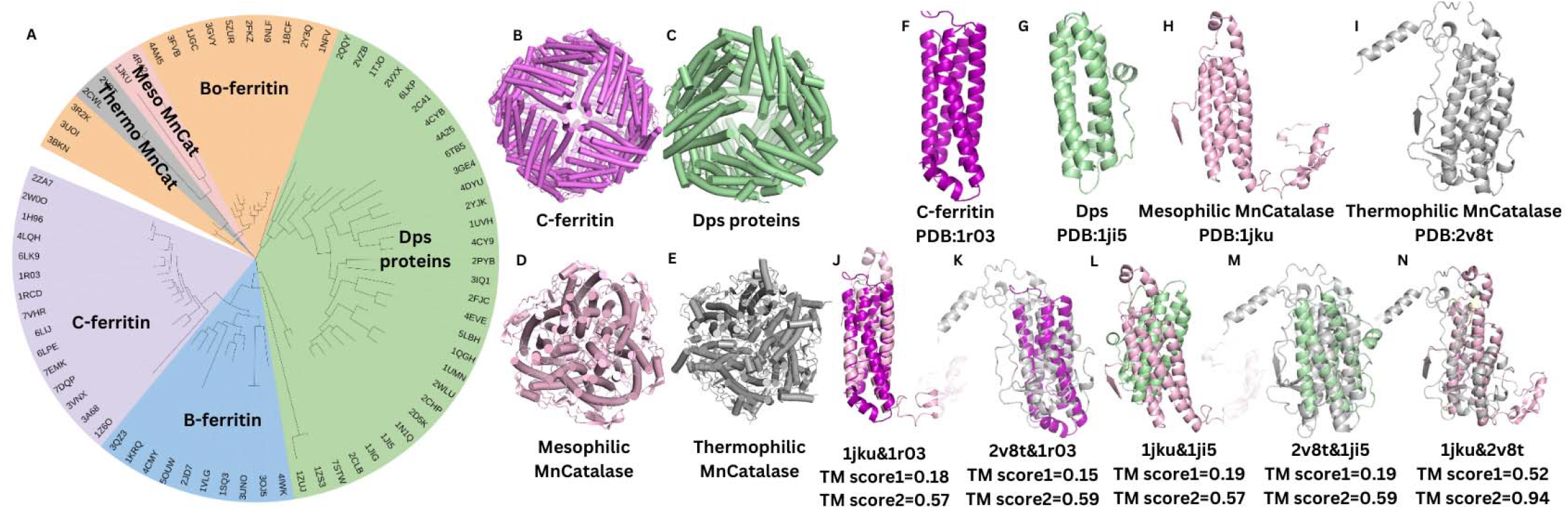
Manganese catalase monomer and its structural attributes. **A.** Phylogenetic Tree of ferritin superfamily in circular layout that includes C-ferritin (purple), B-ferritin (blue), Bo-ferritin (orange), Dps proteins (green), mesophilic manganese catalase (pink) and thermophilic manganese catalase (grey). Cage structure representation of **B**. C-ferritin (PDB:1r03) in purple, **C**. Dps protein (PDB:1JI5) in green, **D**. Mesophilic manganese catalase (PDB:1jku) in pink and **E**. Thermophilic manganese catalase (PDB:2v8t) in grey. **F**. Cartoon representation of Classical ferritin (C-ferritin) subunit comprising 4+1 helices (A.B,C,D helices and a small helix E in purple) (PDB:1r03). **G**. Cartoon representation of Dps protein subunit comprising 4+1 helices (A.B,C,D helices and a small helix E in purple) (PDB:1r03).**H.** Cartoon representation of Mesophilic manganese catalase subunit comprising 4+1 helices (A.B,C,D helices) and a C-terminus extension in pink (PDB:1jku). **I**.Cartoon representation of Thermophilic manganese catalase subunit comprising 4+1 helices (A.B,C,D helices) and a C-terminus extension in grey (PDB:2v8t). Structural alignment between **J**. mesophilic manganese catalase and C-ferritin, between **K**. thermophilic manganese catalase and C-ferritin, **L**. mesophilic manganese catalase and Dps protein, between **M**. thermophilic manganese catalase and Dps proteins and between **N.** mesophilic manganese catalase and thermophilic manganese catalase and their TM scores. TMscore1 depicts the alignment of the full structure, and TMscore2 depicts the alignment of the 4-helix catalytic domain.

#### Secondary structure architecture

Secondary structure composition was analyzed for manganese catalase and ferritin monomers. While α-helix component was most prominent in all the protein types and ⊡-sheet was negligible, distinction was observed in the fraction of ⊡-turns. The fraction of l--turn in mesophilic manganese catalase (f⊡⊡⊡⊡=0.27) and in thermophilic manganese catalase (0.28) is higher than C-ferritin, B-ferritin and Bo-ferritin (0.1) and Dps (0.11) (Table 1, Figure S1 B). This could be attributed to the role of ⊡turns in the folding of helices into compact globular structure (Marcelino and Gierasch, 2008). In case of manganese catalase, compactness and stability conferred by ⊡turns compensate for the long and less ordered C-terminus extensions, thereby stabilizing the monomeric architecture.

#### Amino acid residue composition

We tried to analyze the amino acid composition in manganese catalase proteins with respect to ferritins (Supplementary Table 1). We found mesophilic manganese catalases to have a similar fraction of polar and non-polar residues with respect to ferritin proteins. In contrast, thermophilic manganese catalases were found to have a higher content of non-polar residues than other proteins (Table 1, Figure S1 A). The grand average of hydropathy (GRAVY)(Wilkins et al., 1999) values for proteins were also found to be least negative for thermophilic manganese catalase, validating the observation that thermophilic manganese catalase has a higher content of nonpolar residues. This shift towards more hydrophobic residues contributes to enhanced protein stability and thermostability (Panja et al., 2015).

The molar extinction coefficient was also found to be distinctly higher for manganese catalases than ferritins, suggesting more exposed aromatic residues in catalases than ferritins. However, the absorbance distinction was reduced when normalized with molecular weight. The manganese catalase extensions were rich in amino acid residues A, R/N, G, Q, S, P, E, and K, found prominent in Intrinsically Disordered Regions (IDRs)(Williams et al., 2001). These eight residues constituted more than 55% of the total residue content of manganese catalase extensions, showcasing attributes like the flexibility and polarity of IDRs.

#### Structural alignment study

Structural alignment tools show a weak alignment between manganese catalase and ferritin with a TM-score between 0.17-0.22. However, the structural alignment between only the 4-helix catalytic domain of the two protein groups shows a much stronger alignment with a TM-score above 0.75. The alignment between mesophilic and thermophilic manganese catalase is more potent, with a TM-score of 0.94 for the catalytic domain and 0.52 for the entire structure (Figure 1J-N). This suggests the evolutionary conservation of the catalytic regions between the Manganese catalase and ferritins and divergence regarding the terminal extensions. A similar observation was made across ferritin subfamilies (C-ferritin, B-ferritin, Bo-ferritin and Dps proteins), with terminal extensions conferring structural distinctions and distinct functional attributes to the proteins while conserving the core catalytic properties characteristic of the superfamily(“Physicochemical features of subunit interfaces and their role in self-assembly across the ferritin superfamily,” 2024).

### Characterization of the Subunit Interfaces

#### Interface geometry

Manganese catalase was found to have an interface geometry markedly different from that of the ferritins (Figure 2A). Ferritins are seen to have a 3-chain repetitive unit in their cage assembly comprising an anti-parallel interface (IntA) and two perpendicular interfaces, IntB and IntC (Figure 2B). On the other hand, manganese catalase subunits are arranged sidewise with two parallel side-wise interfaces (we name IntS and IntS’). IntS and IntS’ are distinguished based on the position of the N-terminus extension, whereby the N-terminus is present in IntS and absent in IntS’. The subunits are also angularly aligned such that a chain interacts with the third chain on either side, which we name IntN (Figure 2A). Interface geometry analysis shows mesophilic manganese catalase to have a large buried surface area (BSA) in sidewise interface IntS and thermophilic manganese catalase to have a larger BSA in IntS’. In either case, with manganese catalase, one of the side-wise interfaces is much larger than any of the C-ferritin interfaces. It could be attributed to the long C-terminus extension that wraps around the sideward subunit like a belt, increasing the BSA, Local atomic density (Ld) and thereby enhancing atomic interactions at the interface.

**Figure 2:**
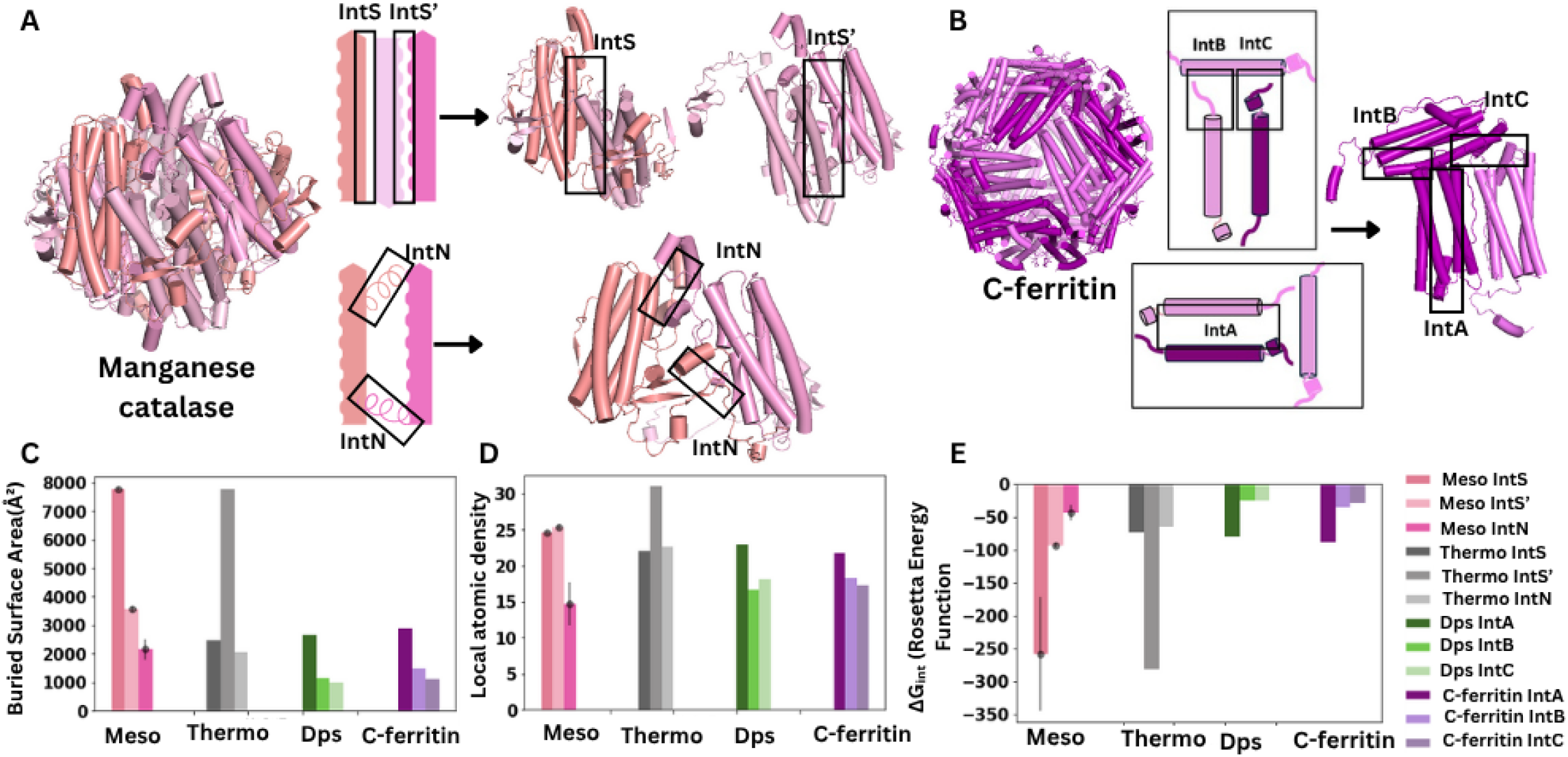
Manganese catalase interface and its structural attributes. **A.** Cylindrical cartoon representation of Manganese catalase (PDB:1jku) showing 3 of the 6 chains comprising sidewise interfaces IntS and IntS’ and angular interface IntN **B**.Cylindrical cartoon representation of C-ferritin (PDB:3a68) showing the cage architecture and 3 of the 24 chains comprising antiparallel interface IntA and perpendicular interfaces IntB and IntC. Distribution of **C**. Buried Surface Area (BSA) and **D.** Local atomic density for the 3 interfaces of Mesophilic (Meso) and Thermophilic (Thermo) Manganese catalase (IntS, IntS’ and IntN) and for the 3 interfaces of C-ferritin and Dps proteins (IntA, IntB and IntC). (Local Atomic Density is the mean number of interface atoms located within 12□ of a given interface atom)(Bahadur et al., 2004) **E.** ΔG analysis for the 3 interfaces of Mesophilic (Meso) and Thermophilic (Thermo) Manganese catalase (IntS, IntS’ and IntN) and for the 3 interfaces of C-ferritin and Dps proteins (IntA, IntB and IntC) using Rosetta tool.In all the barplots, Mesophilic MnCatalase interfaces data is shown in the shades of pink, Thermophilic MnCatalase interfaces data is shown in the shades of grey, C-ferritin interfaces data is shown in the shades of purple and Dps interfaces data is shown in the shades of green

#### Interface electrostatics

The number of H-bond interactions calculated for manganese catalase and ferritin at the interface shows manganese catalase to have a higher count of H-bond interactions for one of the sidewise interfaces as compared to any of the interfaces of C-ferritin (Table 2, Figure S2 A). This property along with larger BSA could have contributed to stabilization the interface with lower ΔG□□□ (Rosetta Energy Function) for the Manganese catalase sidewise interface as compared to the C-ferritin interface. We believe that as Manganese catalase has only 6 subunits in comparison to ferritin, which has 24 subunits, the contribution of each of the manganese catalase interfaces in stabilizing the structure becomes very important. Hence, we observe that one of the side-wise interfaces in manganese catalase has exceptionally high BSA (Figure 2C), Local atomic density (Figure 2D), H-bond number (Figure S2 A) and ΔG□□□ (Figure 2E), helping in stabilizing the interface and for generation of the cage architecture. However, the total contribution of BSA and ΔG for the cage i.e. ΔG□□□ is lower in manganese catalase than in the C-ferritin (Figure S2 B), highlighting the role of greater number of chains and interfaces in stabilizing the quaternary structure.

**Table 2:** Physico-chemical properties of Manganese catalase and C-ferritin interfaces.

|  | Interfaces | Mesophilic<br>MnCatalase(2) | Thermophilic<br>MnCatalase(2) | Interfaces | Dps proteins(30) | C-ferritin(18) |
| --- | --- | --- | --- | --- | --- | --- |
|  | IntS | 7767.8±58.5 | 2450.1±15.2 | IntA | 2646 ± 467 | 2867 ± 330 |
| <Buried surface<br>Area (Å<sup>2</sup>)>, BSA | IntS' | 3575.4±45.9 | 7763.9±106.9 | IntB | 1136± 408 | 1495 ± 511 |
|  | IntN | 2148.8±56.9 | 2061.2±19 | IntC | 1001 ± 358 | 1118 ± 125 |
|  | IntS | 793.7±6.4 | 240.8±1.3 | IntA | 261.1 ± 46.9 | 300.1 ± 36.9 |
| <Number of | IntS' | 359.8±3.9 | 766.7±5.8 | IntB | 120.6 ± 40.9 | 152 ± 47.7 |
| Interface atoms> |  |  |  |  |  |  |
|  | IntN | 215.4±5.2 | 212.1±4.6 | IntC | 105.4 ± 35.4 | 110.9 ± 11.5 |
|  | IntS | 199.9±1.9 | 66±1 | IntA | 71.3 ± 10.6 | 71.2 ± 8.9 |
| <Number of Interface residues> | IntS' | 91±1.2 | 200.7±1.6 | IntB | 31.8 ± 11.2 | 40.2 ± 13.2 |
|  | IntN | 67.1±2.1 | 59.9±0.9 | IntC | 27.3 ± 8.5 | 27.3 ± 8.5 |
|  | IntS | 0.44 | 0.46 | IntA | 0.40 | 0.32 |
| <*f*<sub>non-polar</sub>> | IntS' | 0.61 | 0.43 | IntB | 0.34 | 0.29 |
|  | IntN | 0.38 | 0.33 | IntC | 0.44 | 0.35 |
|  | IntS | 24.6±0.2 | 22±0.1 | IntA | 23.0 ± 1.8 | 21.7 ± 1.7 |
| <Local Atomic Density>Ld | IntS' | 25.3±0.1 | 31±0.3 | IntB | 16.7 ± 2.8 | 18.3 ± 1.5 |
|  | IntN | 14.7±0.4 | 22.7±0.3 | IntC | 18.2 ± 3.0 | 17.2 ± 0.9 |
|  | IntS | 39.2±19.9 | 5.5±1.5 | IntA | 12.6 ± 5.6 | 19.0 ± 6.4 |
| <Number of H-bonds> | IntS' | 17.2±0.7 | 54.5±0.5 | IntB | 5.0 ± 3.7 | 5.6 ± 2.0 |
|  | IntN | 10.5±5.0 | 10.3±0.4 | IntC | 3.1 ± 2.1 | 6.9 ± 1.8 |
|  | IntS | -257.8±86.9 | -72.95±0.65 | IntA | -80±21.9 | -89.1±15.8 |
| <$\Delta G$> (Rosetta energy unit) | IntS' | -92.95±6.65 | -281.55±3.15 | IntB | -24.6±14 | -35.4±13 |
|  | IntN | -43.6±22.7 | -65.85±1.05 | IntC | -25.6±12.2 | -29.1±5.3 |
|  | IntS | 0.76±0.01 | 0.74±0 | IntA | 0.70±0.06 | 0.70±0.04 |
| <Sc> | IntS' | 0.64±0.01 | 0.72±0 | IntB | 0.64±0.06 | 0.67±0.04 |
|  | IntN | 0.65±0.02 | 0.75±0.01 | IntC | 0.72±0.08 | 0.76±0.06 |
|  | IntS | -793 | -950.4 | IntA | -566.6 | -719.8 |
| <Elec> (kcal/mol) | IntS' | -724.2 | -1106.5 | IntB | -539.3 | -667.3 |
|  | IntN | -693.6 | -970.3 | IntC | -520.5 | -662.9 |
|  | IntS | -2975.1, 459.5 | -3582.8, | IntA | -1932.8, | -2162.6, 405.3 |
|  |  |  | 986.9 |  | 535.4 |  |
| <van der Waals energy> attr, rep (kcal/mol) | IntS' | -2780, 559 | -3828.5, 851.6 | IntB | -1885.5, 664.3 | -2110.8, 433.4 |
|  | IntN | -2721.3, 734.1 | -3594.3, 1015.1 | IntC | -1883.3, 759.6 | -2110, 447 |

< $\Delta G$ >: Energy of a biomolecule conformation calculated using REF 15(Alford et al., 2017) in Rosetta energy unit, which includes energies of interactions between non-bonded atom-pairs essential for atomic packing, electrostatics, and solvation; empirical potentials used to model hydrogen- and disulfide-bonds and statistical potentials used to describe backbone and sidechain torsional preferences in proteins.
<Elec>: Electrostatic energy (kcal/mol) calculated using REF 15(Alford et al., 2017)
<van der Waals energy> rep: repulsive energy (kcal/mol); attr: attractive energy (kcal/mol) calculated using REF 15(Alford et al., 2017)
<Sc>: Shape complementarity calculated using REF 15(Alford et al., 2017)
Interface data for C-ferritins and Dps proteins are taken from the author's previous paper("Physicochemical features of subunit interfaces and their role in self-assembly across the ferritin superfamily," 2024)
< $f_{\text{non-polar}}$ >: fraction of non-polar atoms in the interface, averaged over all interfaces of the same type

**Table 3:** Cavity geometry of Manganese catalase, Dps proteins and C-ferritin cages.

|  | **Mesophilic**<br>**MnCatalase** | **Thermophilic**<br>**MnCatalase** | **Dps proteins** | **C-ferritin** |
| --- | --- | --- | --- | --- |
| **Number of chains** | 6 | 6 | 12 | 24 |
| **MW (kDa)** | 170 | 180 | 225 | 475 |
| **Protein diameter**<br>**(angstrom)** | 80 | 80 | 90 | 120 |
| **Protein volume (PV)**<br>**(nm<sup>3</sup>)** | 267 | 270 | 380 | 900 |
| **Cavity volume (CV)(Å<sup>3</sup>)** | 17358 | 17242 | 42915 | 61768 |
| **Cavity surface area**<br>**(CSA)(Å<sup>2</sup>)** | 15353 | 15322 | 31526 | 50931 |
| **Cavity vol/Protein vol**<br>**ratio (CV/PV)** | 0.06 | 0.06 | 0.11 | 0.07 |
| **Cavity surface area /**<br>**Cavity vol ratio**<br>**(CSA/CV) (Å<sup>-1</sup>)** | 0.88 | 0.89 | 0.73 | 0.82 |

We also observe interface electrostatics like van der Waals attraction energy and electrostatic energy higher for thermophilic manganese catalase than mesophilic manganese catalase, while electrostatic properties are similar for mesophilic manganese catalase and C-ferritins (Table2). This data signifies the importance of non covalent interactions in strengthening the subunit interfaces in thermophilic manganese catalases as compared to mesophilic manganese catalase, thereby ensuring the stability of thermophilic manganese catalase at high temperatures.

#### Effect of active-site mutation on the interface geometry and stability

We compared the interface geometry and ΔG□□□ of manganese catalase mutants with that of the wild-type (Mutant PDB: 1o9i, WT: 1jku, Mutant PDB: 4r42, WT: 6j42). We found that mutations at the active site did not affect the interface geometry and ΔG□□□ at the interface of the protein, thus retaining the cage architecture distinctive of manganese catalase (Figure S4).

### Characterization of the Manganese Catalase cavity with respect to Dps proteins and Ferritin

#### Cage geometry

Cavity study of manganese catalase protein shows the cavity volume sum for mesophilic manganese catalase (17358 Å^3^) and thermophilic manganese catalase (17242 Å^3^) as much smaller than that of Dps proteins (42915 Å^3^) and C-ferritins (61768 Å^3^). A similar trend is observed with respect to the sum of cavity surface area with total surface area in mesophilic manganese catalase (15353 Å^2^) and thermophilic manganese catalase (15322 Å^2^) to be lower than Dps proteins (31526 Å^2^) and C-ferritin (50931 Å^2^). However, when we calculated the surface area/volume ratio we found the fraction to be higher in manganese catalase (0.9) than in Dps proteins (0.7) and C-ferritin (0.8) (Figure 3C), explaining a higher surface area exposed to catalysis in manganese catalase than in C-ferritin and Dps protein. Another significant difference is observed in cavity volume to protein volume ratio which is highest for Dps (0.11) and comparable for manganese catalases (0.06) and C-ferritin (0.07). Both these results give us an insight into the cavity architecture disparity between manganese catalase, Dps and C-ferritin. Manganese catalase lacks a central cavity and has multiple small cavities increasing its cavity surface area/ cavity volume ratio, which would contribute to its major function of catalysis. On the other hand, Dps has to accommodate a central cavity like C-ferritin despite only 12 chains in place of 24 chains, hence its cavity volume with respect to protein volume becomes significantly high (0.11) as compared to manganese catalase (0.06) and C-ferritin (0.07). To ensure cage integrity despite the large cavity volume, Dps has ensured more compact structure and low surface area/volume ratio, much less than manganese catalases and even less than C-ferritin.

**Figure 3:**
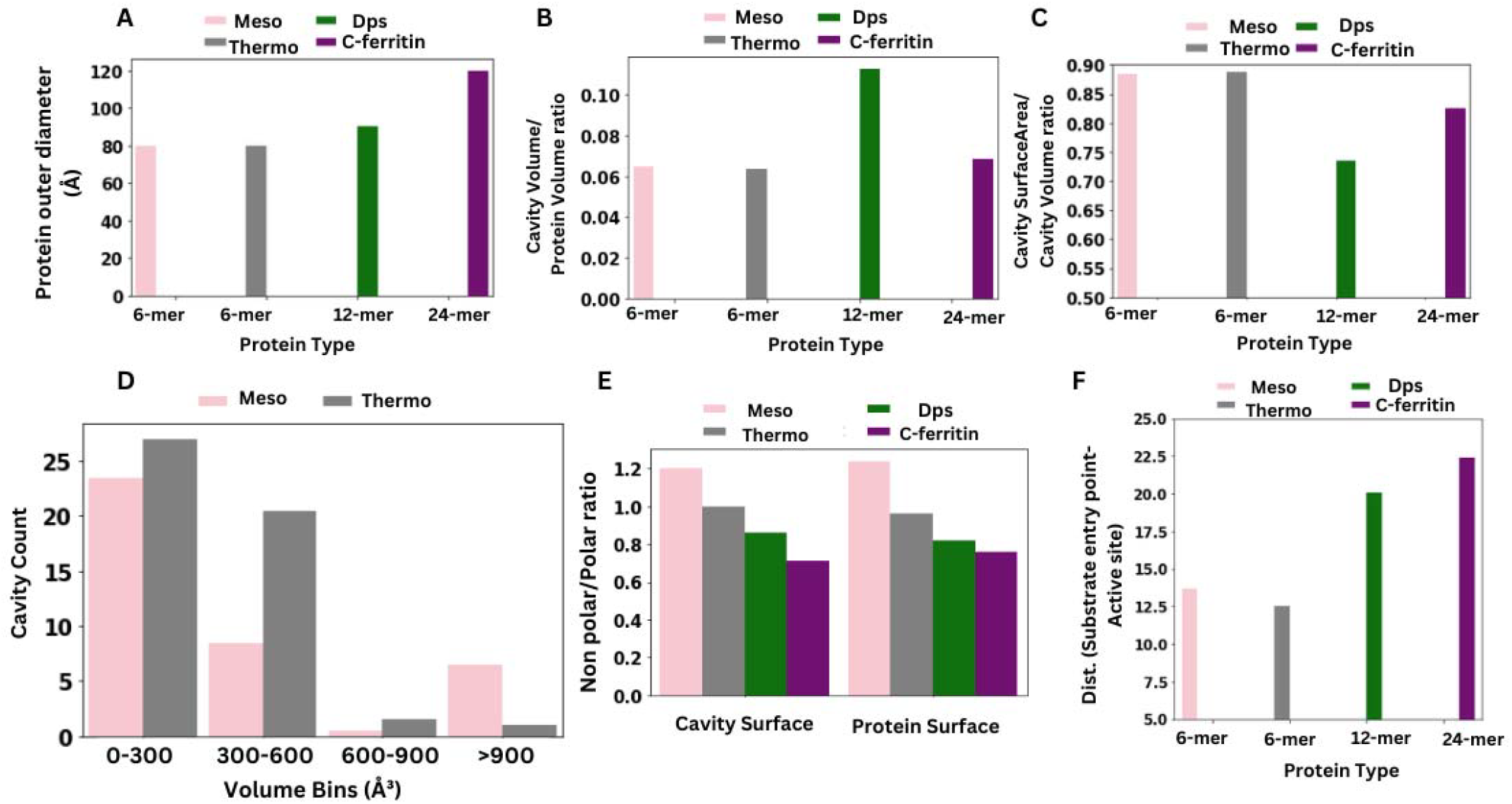
Manganese catalase cavity with respect to Dps proteins and C-ferritin. **A**. Barplot depicting outer cage diameter in Å of Mesophilic Manganese catalase in pink, Thermophilic Manganese catalase in grey, Dps proteins in green and C-ferritin in purple. **B**. Barplot depicting Cavity volume by Protein volume ratio using the same color code as in A **C**. Barplot depicting Cavity surface area by cavity volume using the same color code as in A **D**. Frequency distribution bars showing the cavity count for different volume bins. **E**. Barplot depicting Non-polar residues to polar residues ratio for cavity surface and protein surface for proteins color coded as in A and, **F**. Distance between metal entry point and catalytic site in Å for proteins color coded as in A.

#### Cage polarity

Polarity of the cavity surface with respect to protein surface was analyzed for manganese catalase and C-ferritin and a clear distinction was made between the two proteins. It was observed that the ratio of non polar/polar amino acid residues in cavity surface was higher for mesophilic manganese catalase (NP/P ratio = 1.2) and thermophilic manganese catalase (NP/P ratio = 1.0) than the C-ferritin (NP/P ratio = 0.7). Similar trend was observed for protein surface whereby mesophilic manganese catalase (NP/P ratio = 1.24) and thermophilic manganese catalase (NP/P ratio = 0.96) showed higher Non-Polar/ Polar amino acid residues than the C-ferritin (NP/P ratio = 0.76). This trend confirms the hydrophilic properties of ferritin interior and exterior with respect to Manganese Catalase.The hydrophilic interior of ferritin is known to pose a challenge to the delivery of hydrophobic drugs by ferritin based drug delivery system(Incocciati et al., 2023). On the contrary, the hydrophobic environment in manganese catalases ensures fast diffusion of polar substrates contributing to the rapid catalytic activity observed in catalases (Zámocký and Koller, 1999). Between the two manganese catalase subtypes, mesophilic manganese catalase cavities have more hydrophobic character than thermophilic manganese catalase. However, the higher polarity of C-ferritin surface depicted by lower NP/P ratio confers higher solubility to ferritin as compared to both mesophilic and thermophilic manganese catalase(Ptak-Kaczor et al., 2021). In ferritin, the active site is the ferroxidase centre where the rapid oxidation of Fe(II) to Fe(III) occurs. The location of this site plays a critical role in iron sequestration and iron storage(Ciambellotti et al., 2021). We found ferritin catalytic site to be located at a greater distance from the ion entry point (C-ferritin=22.4 Å) than that of manganese catalase (Mesophilic MnCatalase=13.7Å and Thermophilic MnCatalase=12.5 Å). This could be attributed to the small size of the manganese catalase cage as well as the need for manganese catalase active site to be closer to the ion entry point for its catalytic activity to be efficient given that manganese catalase has one of the highest turnover rates among all enzymes (Smejkal and Kakumanu, 2019).

## Methods

### Dataset

RCSB PDB was used to collect initial information on the deposited structures of manganese catalase. BLAST was undertaken to obtain homologous sequences of mesophilic and thermophilic manganese catalases using PDB: 1jku and 2v8t as reference sequences and selection criteria of >60% sequence identity and >80% coverage, taking the top 30 hits in our dataset and also ensuring organism-wise nonredundancy. Ferritin datasets were taken from our previous study(“Physicochemical features of subunit interfaces and their role in self-assembly across the ferritin superfamily,” 2024) .

*Monomer sequence characteristics– GRAVY, Secondary structure fraction, Instability Index* The physicochemical properties of manganese catalase and ferritin monomers were studied using the Protparam tool (Wilkins et al., 1999). Amino acid composition, aliphatic index, instability index and grand average of hydropathicity (GRAVY) were generated to compare these properties at a monomer scale for manganese catalases, Dps proteins and Classical ferritins. Monomer secondary structure composition was analyzed using PDB structures and DSSP tool(Kabsch and Sander, 1983).

### Interface characteristics– area, packing, geometry

Each protein was split into subunits, and the Buried Surface Area (BSA) was estimated as the sum of Solvent Accessible Surface Area (ASA) of the two subunits, less that of the pair (i.e the complex form). Residues/atoms that lose ASA after complexation are considered to be interface residues/atoms. ASA calculation was done using the program NACCESS(Hubbard, 1993), which implements the Lee and Richards algorithm(Lee and Richards, 1971). The atomic packing at the interface was done by calculating the local atomic density L_D_ (Bahadur et al., 2004). The interfaces were clustered according to their size (BSA) and then by manual inspection (using PYMOL), we categorized them into three types based on their size and geometry of interaction – a) sidewise interfaces (IntS and IntS’), b) angular interface (IntN). For C-ferritin and Dps, the interfaces are categorized as a) anti-parallel interfaces (IntA), b) perpendicular interface with N-terminus extension (IntB), and c) perpendicular interface with E-helix and C-terminus extension (IntC).

### Interface stability

We used Rosetta (Alford et al., 2017), an all atom energy function of macromolecules for calculating all the energy components and the overall free energy of binding/ energy of a biomolecule (ΔG) for each of the interface types of all ferritin assemblies. The energy unit using REF 15 is described in “Rosetta Energy Unit”, as in addition to molecular mechanics energy terms which includes energies of interactions between non-bonded atom-pairs important for atomic packing, electrostatics, and solvation. Rosetta also provides empirical potentials including van der waals potential and electrostatic potentials to understand the energetics of interface interactions. All the molecular mechanics energy terms are estimated in kcals/mol.

### Cage characteristics– cavity volume, cavity surface area, geometry, surface polarity

CavityPlus web server (Xu et al., 2018) was used for the generation of cavity files of manganese catalase, Dps and C-ferritin cages. These cavity files were used for obtaining cavity volume, cavity surface area and cavity surface information. These were utilized to generate cavity surface area by volume ratio, cavity volume by protein volume ratio and cavity polarity. Protein surface polarity was obtained from the Accessible Surface Area (ASA) files of the protein cage obtained from the NACCESS (Hubbard, 1993). Manganese Catalases, Dps and ferritin volumes are obtained from the literature.

## Conclusion

Our study tries to give structural answers to the functional properties observed in manganese catalases with respect to Dps and C-ferritins. We found distinct subunit-subunit interfaces in manganese catalase and greater interface area, compactness and ΔG□□□ per interface with respect to Dps proteins and Classical ferritins that makes cage architecture possible in them despite the presence of only 6 chains. We also observed differences in thermophilic and mesophilic manganese catalases with the thermophilic manganese catalases having higher number of non-polar residues at the monomer level and, greater electrostatic and van der Waals interactions at the interface level, possibly contributing towards greater thermo-chemical stability in them. Our study also gives insight into cage architecture of manganese catalases with respect to Dps and Classical ferritins, whereby greater surface area/volume ratio of the cage cavity, higher propensity of non-polar residues at the cavity surface as well as smaller distance between the substrate entry point and active site could be expected to confer high catalytic efficiency to manganese catalases. We also observe a transition from no central cavity in manganese catalase to the presence of central cavity in Dps proteins and C-ferritins. To accommodate a central cavity despite 12 chains, Dps reduces the cavity surface area/ cavity volume parameter (less than manganese catalase and even C-ferritins) to ensure higher compactness and sustenance of the cage despite the large central cavity. Thus, this study offers an insight into structural variations in the interfaces and cage architecture of manganese catalases, Dps and C-ferritins giving an insight into the transition of their function from catalysis to storage activities .

## Author Contributions

SD and SC have conceived the idea. A has done all the experiments and analyses. A and SD wrote the manuscript with input from all authors.

## Acknowledgments

A acknowledges the fellowship from CSIR-NET India. SD acknowledges DBT India for the RLS grant (RLS grant to SD: BT/ RLF/Re-entry/10/2020, sanction order serial number 145) and IIT Jodhpur India for infrastructure support.

