## Supplementary information for "Comparative study of manganese catalase monomers, interfaces and cage architecture within the ferritin superfamily"

**Supplementary Table 1.** Amino acid composition in a single (monomeric) unit of manganese catalase proteins and other ferritin monomers

| | Mesophilic MnCatalase | Thermophilic MnCatalase | C-ferritin | B-ferritin | Bo-ferritin | Dps proteins |
|---|---|---|---|---|---|---|
| ALA | 21 | 31 | 15 | 17 | 13 | 17 |
| CYS | 0 | 0 | 2 | 1 | 1 | 1 |
| ASP | 17 | 16 | 12 | 11 | 12 | 11 |
| GLU | 24 | 23 | 18 | 18 | 19 | 16 |
| PHE | 9 | 14 | 8 | 9 | 4 | 6 |
| GLY | 25 | 27 | 9 | 7 | 10 | 9 |
| HIS | 8 | 9 | 7 | 8 | 7 | 7 |
| ILE | 7 | 14 | 6 | 8 | 10 | 9 |
| LYS | 15 | 16 | 12 | 9 | 9 | 9 |
| LEU | 24 | 32 | 20 | 19 | 22 | 18 |
| MET | 15 | 11 | 5 | 6 | 5 | 4 |

| | | | | | | |
|---|---|---|---|---|---|---|
| ASN | 10 | 14 | 8 | 7 | 6 | 6 |
| PRO | 12 | 19 | 8 | 7 | 6 | 6 |
| GLN | 13 | 7 | 10 | 9 | 9 | 7 |
| ARG | 12 | 13 | 8 | 7 | 8 | 7 |
| SER | 15 | 12 | 11 | 11 | 6 | 8 |
| THR | 14 | 10 | 6 | 7 | 6 | 10 |
| VAL | 12 | 16 | 10 | 9 | 6 | 11 |
| TRP | 3 | 2 | 1 | 2 | 2 | 2 |
| TYR | 9 | 13 | 8 | 6 | 6 | 6 |

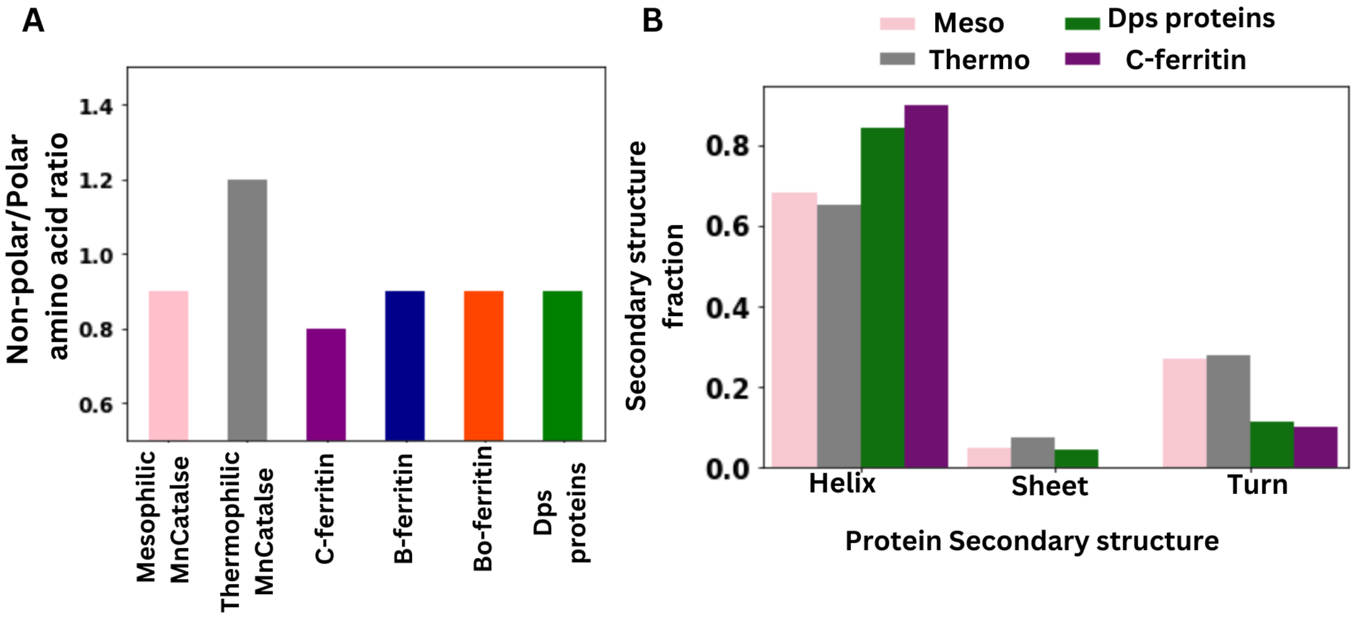


**Figure S1: Comparison of Non-polar/Polar amino acid ratio and Secondary structure fraction between Manganese Catalases and Classical ferritins (C-ferritin), Bacterial ferritins (B-ferritin), Bacterioferritin (Bo-ferritin) and Dps proteins**

**A.** Non-polar to polar amino acid ratio following the color code as for Mesophilic manganese catalase (pink), Thermophilic manganese catalase (grey), C-ferritin (purple), B-ferritin (blue), Bo-ferritin (orange) and Dps proteins (green)**B.** Distribution of Secondary structure fraction following the color code as for Mesophilic manganese catalase (pink), Thermophilic manganese catalase (grey), C-ferritin (purple), B-ferritin (blue), Bo-ferritin (orange) and Dps proteins (green).

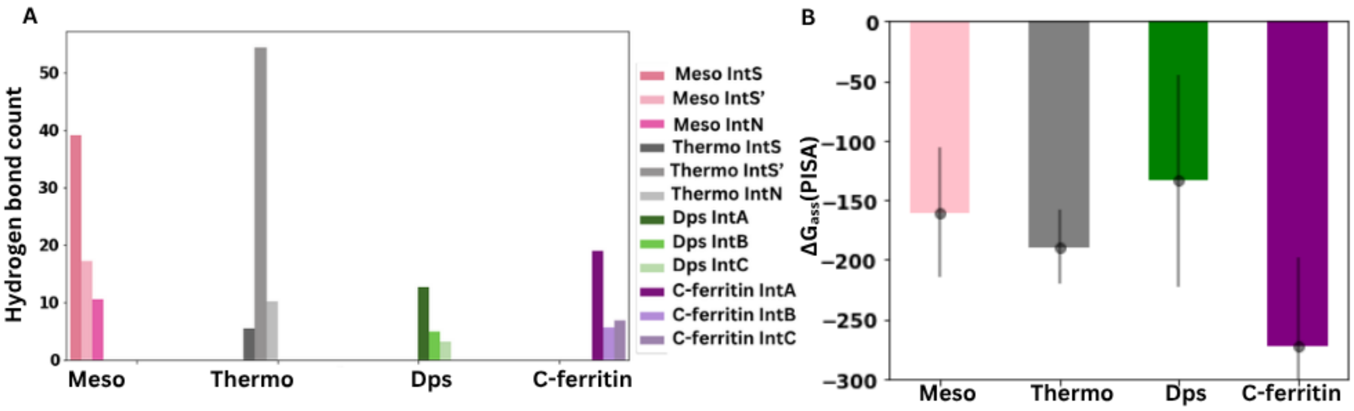


**Figure S2: Comparison of H-bond count and $\Delta G_{ass}$ between Manganese Catalases and C-ferritin**

Barplot of **A**. H-bond count at interfaces of Mesophilic Manganese catalase in pink, Thermophilic Manganese catalase in grey and C-ferritin in purple **B**. $\Delta G_{ass}$ of Mesophilic Manganese catalase in pink, Thermophilic Manganese catalase in grey, C-ferritin in purple and Dps in green as obtained from PISA

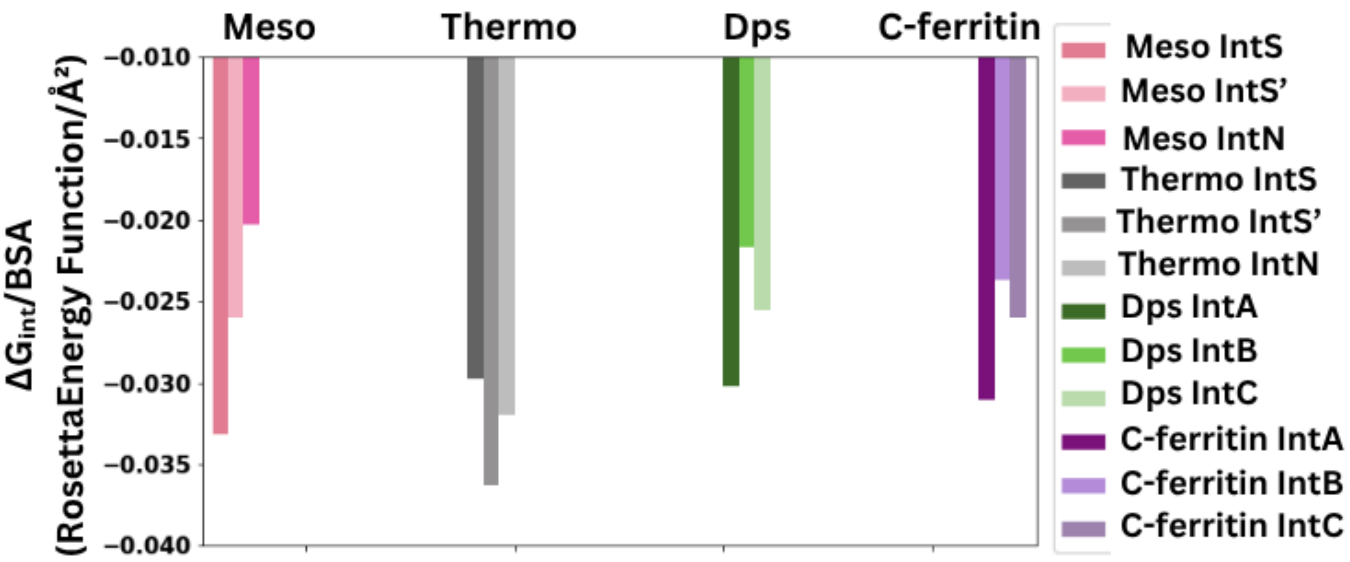


**Figure S3: Comparison of $\Delta G_{int}$/BSA between Manganese Catalases, C-ferritin and Dps proteins**

Barplot of $\Delta G_{int}$/BSA at interfaces of Mesophilic Manganese catalase in pink, Thermophilic Manganese catalase in grey and C-ferritin in purple and Dps in green

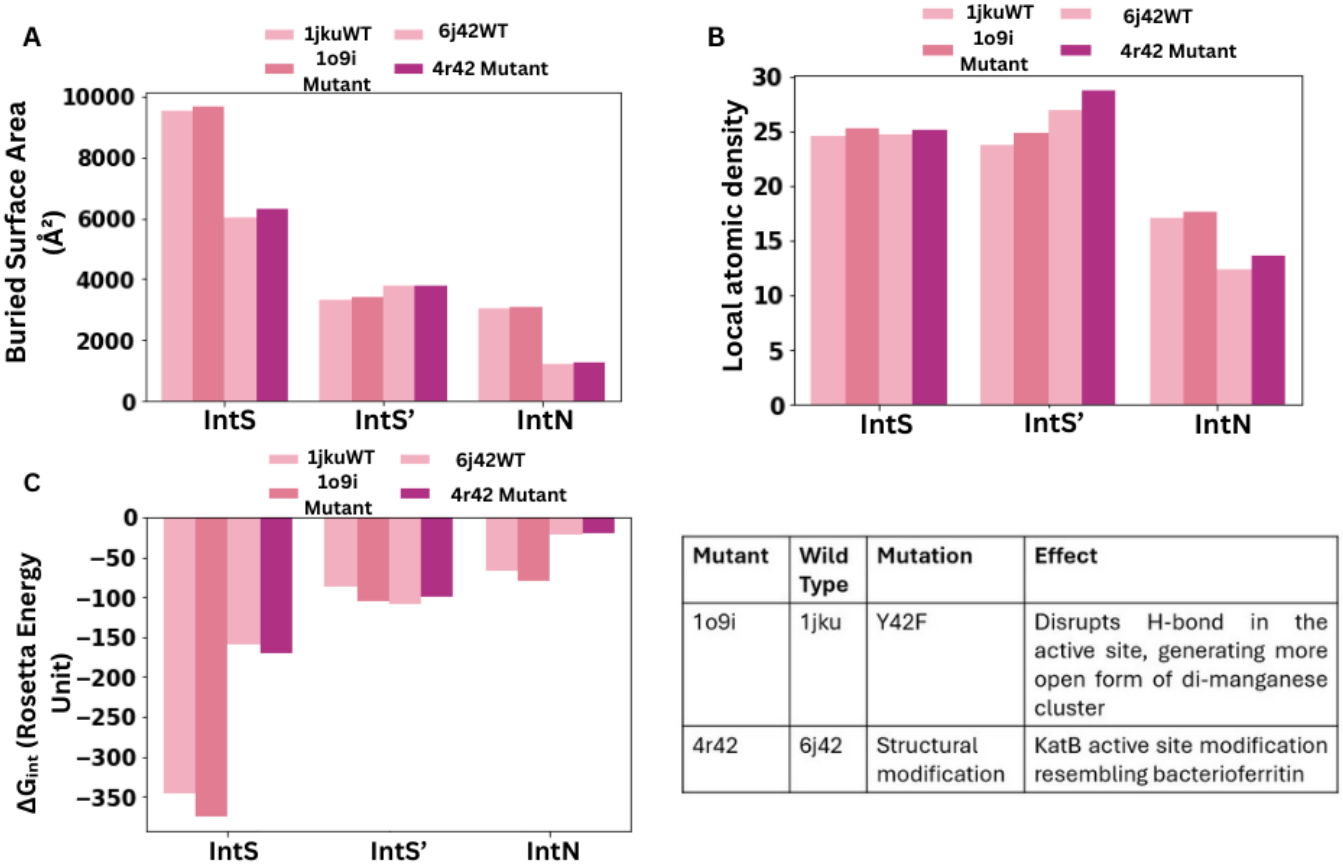


| Mutant | Wild Type | Mutation | Effect |
|---|---|---|---|
| 1o9i | 1jku | Y42F | Disrupts H-bond in the active site, generating more open form of di-manganese cluster |
| 4r42 | 6j42 | Structural modification | KatB active site modification resembling bacterioferritin |

**Figure S4: Comparison of A. BSA, Local atomic density and $\Delta G_{int}$ between Wild Type and mutant Manganese catalase**

Barplot of **A**. BSA **B**.Local atomic density and **C**.$\Delta G_{int}$ at interfaces IntS, IntS' and IntN for wildtype Manganese catalase (PDB:1jku, 6j42) depicted in lighter shade of pink and mutant Manganese catalase (PDB:1o9i, 4r42) depicted in darker shade of pink
